# Wireless *in vivo* Recording of Cortical Activity by an Ion-Sensitive Field Effect Transistor

**DOI:** 10.1101/2023.01.19.524785

**Authors:** Suyash Bhatt, Emily Masterson, Tianxiang Zhu, Jenna Eizadi, Judy George, Nesya Graupe, Adam Vareberg, Jack Phillips, Ilhan Bok, Matthew Dwyer, Alireza Ashtiani, Aviad Hai

## Abstract

Wireless brain technologies are empowering basic neuroscience and clinical neurology by offering new platforms that minimize invasiveness and refine possibilities during electrophysiological recording and stimulation. Despite their advantages, most systems require on-board power supply and sizeable transmission circuitry, enforcing a lower bound for miniaturization. Designing new minimalistic architectures that can efficiently sense neurophysiological events will open the door to standalone microscale sensors and minimally invasive delivery of multiple sensors. Here we present a circuit for sensing ionic fluctuations in the brain by an ion-sensitive field effect transistor that detunes a single radiofrequency resonator in parallel. We establish sensitivity of the sensor by electromagnetic analysis and quantify response to ionic fluctuations *in vitro*. We validate this new architecture *in vivo* during hindpaw stimulation in rodents and verify correlation with local field potential recordings. This new approach can be implemented as an integrated circuit for wireless *in situ* recording of brain electrophysiology.

## 1. INTRODUCTION

Wired probes are currently the mainstay devices for *in situ* recording and stimulation of brain tissue[1–4]. However, wireless variants are becoming ubiquitous in both neurobiological research[5–8] and neurological applications[9–11], greatly expanding the spectrum of behaviors and brain functions studied. Most wireless electrophysiology systems are reliant on an on-board battery for data transmission or include circuitry for power harvesting, employing either head-mounted replaceable power cells[7,6,12] or advanced transcutaneous inductive links for periodical charging and continuous operation[5,8]. More minimalistic circuits that interact with outside detectors such as ultrasound transducers[13,14] magnetic resonance imaging (MRI) hardware[15– 18] and optoelectronic interfaces[19–22] present new possibilities for backscatter power harvesting, simplified device delivery and localization, and detection across multiple regions in the nervous system. Radio frequency (RF) sensors in particular, have traditionally benefitted from high signal penetration through the skull and other tissue types compared with optical and ultrasonic modalities[23,24], and new designs enable improved deep tissue sensing, micro-scale device localization, and controlled drug delivery[25–28]. Recently, tunable RF sensors were implanted cortically and used for wireless, MRI-mediated *in vivo* monitoring of bioluminescent cell grafts in the brain[15]. By using field effect transistors (FETs) as a tuning element that modulates frequency response of implanted resonating antennae connected in parallel to the FET, versatile biophysical and biochemical cues in the brain and other organs can be sensed wirelessly and relayed to outside RF detectors[15,16]. The integration of ion-sensitive FETS (ISFETs) with tunable RF resonators has yet to be demonstrated but offers an enticing possibility for wireless rapid detection of ionic fluctuations in the brain related to stimulation-evoked activity and the analysis of different brain states.

ISFETs have been used as wired probes to directly detect neuronal activity from isolated cells[29–31], brain slices[32,33] and for neurotransmitter monitoring via chemical functionalization[34–36]. Relying on modified fabrication processes, the FET gate terminal is exposed to electrolyte, and acts as a micro- or nano-scale sensing electrode responsive to biophysical events that induce a change in FET channel transconductance[37,38]. This same mechanism can be leveraged to detune a resonator, similarly to photosensitive FETs used to detune wireless resonators in response to light[15,39]. In a similar topology, an ISFET would act as an electric shunt, diverting current away from the resonating element in response to ionic events. When coupled to RF detector inductively, such a device could modulate coupling between ISFETs implanted *in situ* and detection hardware and therefore facilitate readouts specific to changes in the ionic makeup occurring at the active microenvironment of the device (Fig. 1).

**Fig. 1.**
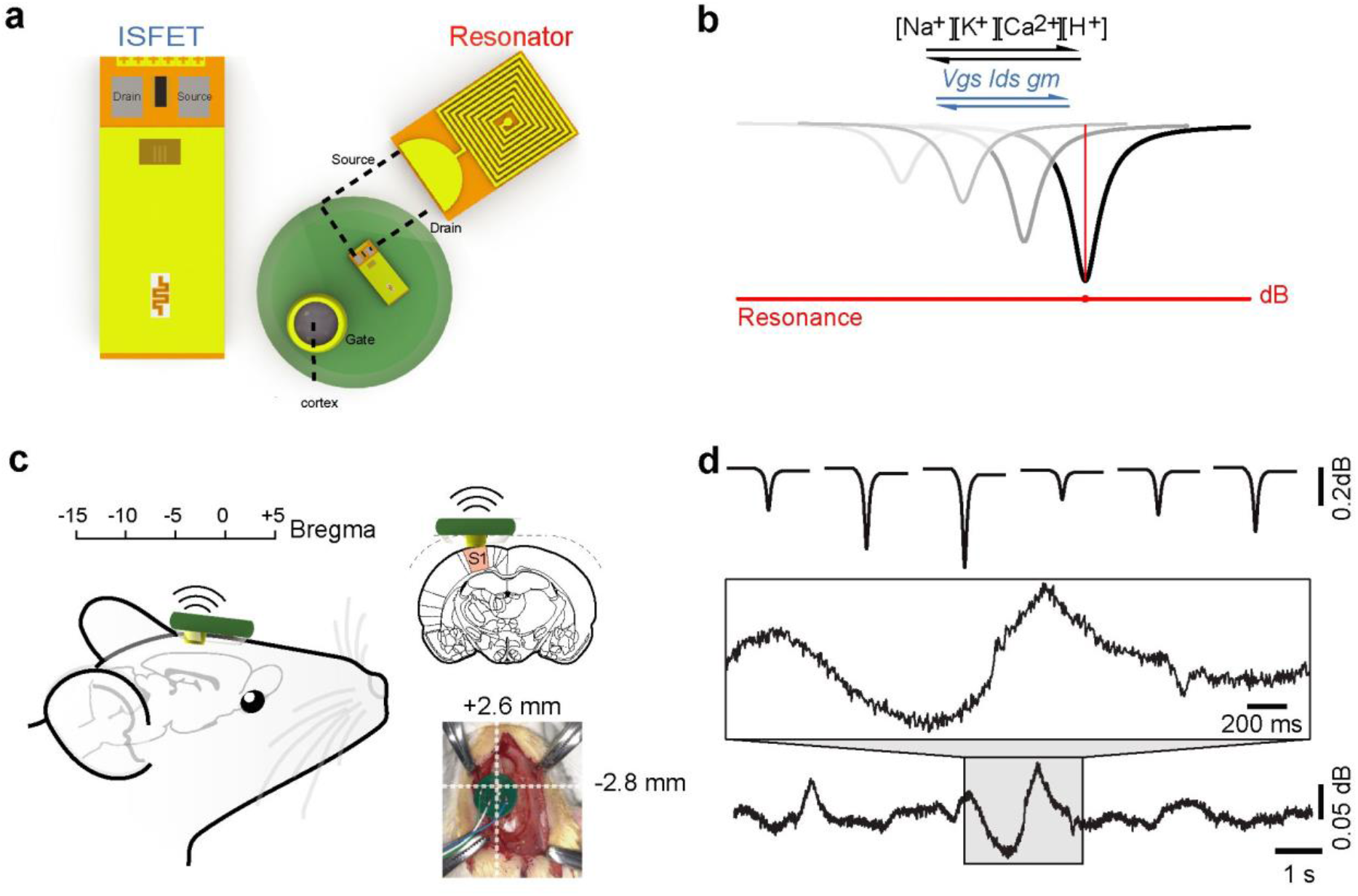
Wireless ion sensitive field effect transistor (ISFET) for *in vivo* cortical recordings of ionic fluctuations. **a** Source and drain of ISFET device are connected to top and bottom plates of capacitor, linking in parallel to circuit. **b** Q of resonator is dependent on ionic concentrations local to ISFET gate electrode. **c** Active site of ISFET is embedded through cranial window on surface of somatosensory cortex. **d** Ion fluctuations detected wirelessly represented in time domain by S11 minima between resonator and antenna over 60 s window.

The intent of this study is to demonstrate the validity of this approach. We present a circuit for sensing ionic fluctuations by an ISFET device that detunes an RF resonator connected in parallel to the FET drain and source terminals (Fig. 1a). We make computational predictions of the sensitivity of the circuit by way of a combined semiconductor and electromagnetic simulation environment. We then validate our predictions by wireless measurement of ionic levels in physiological solution. Finally, we demonstrate the use of this new architecture by an implanted ISFET connected to a resonator in the cortex of anesthetized rats and perform wireless frequency response measurements during hindpaw stimulation. Using wired microelectrode arrays during the same stimulation paradigm, we correlate between the frequency response and local field potential amplitudes for a repertoire of responses corresponding to positive and negative phase events. This recording technique can be a basis for miniaturized devices comprising resonators and ISFETs integrated on the same silicon chip and used for wireless recording of brain electrophysiology.

## 2. METHODS

### 2.1 Electromagnetic simulations

Simulations of device response were performed using a combination of Silvaco TCAD 4.6.2.R software suite (Silvaco Inc., Santa Clara, CA) for ISFET characterization, and Sonnet 17.52 Electromagnetic Software (Sonnet Software, Syracuse, NY) for radio frequency (RF) device detuning simulations. ISFET S-parameters were exported from Silvaco and input into Sonnet using in-house code available through GitHub (see Supplementary Information). ISFET fabrication process was emulated in two dimensions via Silvaco Athena module, whereby an SOI wafer (Si/SiO_2_/Si, 0.8/3/0.2 µm) was selectively doped with Boron to form the P-well at a dose of 8.5·10^12^/cm^2^ with an energy of 100 keV diffused using the Pearson method (diffusion time was 310 sec at 1200 °C). A layer of nitride (50 nm) was used as a sacrificial barrier to block the dose at the source and drain. An additional barrier dose of Boron was implanted into the channel at 1·10^11^/cm^2^ with energy of 10 keV. Following removal of the nitride barrier layer, a layer of polysilicon was deposited to protect the p-well, and a dose of arsenic at 5·10^15^/cm^2^ with an energy of 50 keV was deposited at the source and drain terminals. The dose was diffused at 900 °C for 2 sec, followed by etching of the polysilicon sacrificial layer. The electrolyte solution facing the active site of the ISFET was modeled as a modified germanium layer (1 µm x 1 µm) for default properties, with relative permittivity (80), charge carrier mobility (µ_n_ = 6.9·10^−4^ cm^2^/(V·s), µ_n_ = 5·10^−4^ cm^2^/(V·s)), and electron affinity (3.9 eV) modified based on previous work to emulate an electrolyte solution[40]. A gold reference electrode (1 µm x 0.1 µm) was then deposited above electrolyte layer, enabling forward voltage bias to the gate and emulating ISFET sensing. The large signal model of the ISFET was extracted by fixing *V*_*ds*_ at 5 V and sweeping *V*_*gs*_ from 0 V to 3 V with steps of 0.1 V. The transconductance *g*_*m*_ was quantified by taking the derivative of the current as a function of *V*_*gs*_ from the large signal model, modeling the small signal gain of the ISFET. ISFET S-parameters were extracted from the small signal model for coupling to opposing sides of an inductor-capacitor (LC) resonator modelled in Sonnet. The resonator comprised a 10-turn square inductor (W/L: 3.5/3.5 mm) and a 30 × 50 mm parallel plate capacitor with turn width of 75 µm and turn spacing of 75 µm, and metal layer thickness of 36 µm. The device was coupled to a model antenna with 1 mm distance to simulate the near field coupling representative of the *in vivo* mode, as well as the induced magnetic field *B*. The antenna was a single turn spiral with an outer diameter of 9 mm, and inner diameter of 2.5 mm. Single port S11 coupling coefficient was calculated as a function of both frequency and *V*_*gs*_ corresponding to ionic concentrations proximal to the gate. Frequency response sweeps were simulated from 100 to 200 MHz with a step size of 500 kHz and tested with a corresponding pH sweep of 6 to 8 with intervals of 0.5 (equivalent to 17.5 mV).

### 2.2 Frequency response measurements

High speed measurements of device frequency response were performed using Copper Mountain 1-port R60 vector network analyzer (VNA) (Copper Mountain Technologies, Indianapolis, IN) or Keysight E5061A (Keysight Technologies, Santa Rosa, CA) and controlled by in-house python scripts (Supplementary Information). S11 frequency sweeps were read via SMA-connected PCB transceiver antenna (RFEAH-5, RF Explorer Technologies, Madrid, Spain) and controlled via USB or TCP/IP port connection based on existing VNA API. Data was acquired via binary bin-block transfer at sweep rates of up to 145 Hz at a range of 51 to 201 datapoints and corresponding timestamps per sweep. For both *in vitro* and *in vivo* experiments, the drain and source terminals of a MSFET 3351 ion sensitive field effect transistor (ISFET) (Microsens SA, Lausanne, Switzerland) were bonded in parallel to a standard printed-circuit resonator comprising gold-plated copper (thickness of 36 µm) ten-turn square inductors (W/L: 3.5 × 3.5 mm, turn width: 75 µm, turn spacing: 75 µm) and a 30 × 50 mm parallel plate capacitor.For *in vitro* frequency response measurements, an average of 10 sweeps was taken for each test solution titrated at pH levels ranging between 6.5 and 7.5. The sensing gate terminal of the ISFET was immersed in test solution and allowed 400 ms to reach steady state before frequency dependent S11 coupling coefficient was recorded.

### 2.3 Animal use

Sprague-Dawley rats 8-10 weeks of age (250g) were purchased from Envigo (Madison, WI) and used for all *in vivo* experiments. The animals were housed and maintained on a 12 h light/dark cycle and permitted ad libitum access to food and water. All procedures were performed in strict compliance with US Federal guidelines, with veterinary oversight and approval by the UW-Madison Research Animals Resource Center (RARC).

### 2.4 Surgical procedures

In preparation of electrophysiological recording experiments, rats were anesthetized (isoflurane, 4% induction, 1.5% maintenance) shaved and mounted on a stereotaxic device (Harvard Apparatus, Holliston, Massachusetts). Heart rate and blood oxygenation were continuously monitored using a Nonin 8600 series pulse oximeter (Nonin Inc., Plymouth, MN) and animals were secured with ear bars during all subsequent procedures. Heart rates were maintained at 360-380 beats per minute. The scalp was retracted and holes were drilled into the skull using 1.6 mm diameter drill bits (Cellpoint Scientific Inc., Gaithersburg, MD) with coordinates of 2.2 mm posterior and 2.0 mm lateral to bregma into the somatosensory cortex (trunk field) region. For ISFET recordings, a cranial window of 3 mm diameter was exposed by drilling several adjacent holes with 1.6mm drill bit. Microelectrode arrays (MEAs) or ISFETs were attached to a stereotaxic arm and manipulated into exposed cortical region (Fig. 1). The microelectrode array was lowered into the somatosensory cortex 1.5 mm below the dura and swept at depths ranging between 2 and 4 mm. The reference electrode was placed into tissue around the skull and the stereotactic frame was grounded to minimize noise. For hindpaw stimulation during brain recording, a stimulating electrode (part #: ELSTM1, BIOPAC Systems, Inc., Goleta, California) was inserted through the right contralateral hindpaw.

### 2.5 In vivo recording and stimulation

Microelectrode array (MEA) recordings were performed using 4-channel Q1×1-tet-5mm-121-CQ4 and 16-channel A1×16-5mm-25-177-A16 wired probes (Neuronexus, Ann Arbor, Michigan) fed into a low profile non-ferrous headstage (LP16CH-ZNF) and streamed through a PZ5-64 neurodigitizer amplifier and RZ2-2 base processor (Tucker-Davis Technologies, Alachua, FL). Raw data were acquired at 65 MHz sampling rate per channel and loaded into MATLAB R2021a (Mathworks, Natick, MA) using a customized TDT software development kit and in-house routines for processing and analysis. Electrical hindpaw stimulation was applied with 1 msec 5 mA current pulses using an isolated current stimulator (A382, World Precision Instruments, Sarasota, Florida). For each stimulation condition, a 60 second epoch was recorded beginning with a 15 sec pre-stimulation baseline, followed by a 30 sec stimulation period at frequencies of 2, 5, or 10 Hz and concluding with a 15 second baseline immediately after stimulation offset. For each animal, three randomized trials were conducted for each stimulation condition. High speed frequency response of brain-implanted ISFET resonator was acquired before, during and after hindpaw stimulation with identical parameters, at sampling rate of 145 Hz by SMA-connected near-field antenna (RFEAH-5, RF Explorer Technologies, Madrid, Spain).

### 2.6 Data analysis

Amplitude of device frequency response during hindpaw stimulation was quantified by first taking the on-resonance S11 minima for each trace and calculating a baseline root mean square (RMS) along two random 2 sec segments of inactivity during the 60 sec baseline recording prior to stimulation. Response amplitude was then defined as the RMS during a 2 sec window immediately upon onset of stimulation divided by the baseline RMS value of the same experiment. Frequency domain beta band analysis was done by generating the power spectrum of the first 5 seconds following onset of stimulation normalized to the power spectrum of the 15 seconds before onset of stimulation to account for drift of the ISFET and differences in impedances of the electrode from trial to trial. LFP frequency band responses of corresponding stimulation rate were analyzed individually to determine correlation post onset of stimulation.

## 3. RESULTS

### 3.1 Electromagnetic modeling of device performance

To predict the sensitivity of the device and its ability to generate wireless readouts of physiological ionic fluctuations and pH levels, we used finite element electromagnetic modeling in conjunction with semiconductor simulations to mimic the ISFET-coupled resonator (Fig. 2). The resonator coupled with the antenna was modeled to reflect the induced electric field at different impedance values of the ISFET channel surrounding resonance (Fig. 2b-c). We estimate a Q factor of 127 at initial tuning of the device corresponding to maximum e-field at resonance of 1.4 × 10^3^ V/m (Fig. 2b). Modulation of Q factor ranged between 63.5 and 127 for corresponding ISFET gate channel impedance values ranging between 4500 and 500 Ohm corresponding to input *V*_*ov*_ values ranging between 482.5 and 500 mV and pH levels ranging between 6.5 and 7 (Fig. 2c). The modeled ISFET (Fig. 2d) was designed with a doping profile that most closely resembles the threshold voltage of the ISFET used for *in vivo* experiments and displayed a relatively deep channel (0.2 µm) where current was generated at a density of 4.84 × 10^−7^ A/cm^2^ with a modest bias voltage of 0.5V and *V*_*ds*_ of 3V (Fig. 2e-f) shown previously to be induced by coupled antennae during frequency sweep at the MHz regime[15]. Input *V*_*gs*_ sweep revealed linear current response at 0.5 V overdrive voltage (Fig. 2e) with a transconductance of approximately 0.8 mS (Fig. 2f). To obtain the expected response to physiological ionic concentrations, a 10 µm gate channel ISFET model displaying characteristic pH response (35 mV/pH) was combined with the RF antenna (Fig. 2, a and g). The frequency response at pH levels ranging between 6 and 8 showed Q factor modulation ranging between 18.1 and 42.3 and average gain modulation of −0.08 dB/pH (Fig. 2g, inset) suggesting detectability to changes in ion concentrations of as low as .068 µM well below changes commonly seen in the brain interstitium[41,42].

**Fig. 2.**
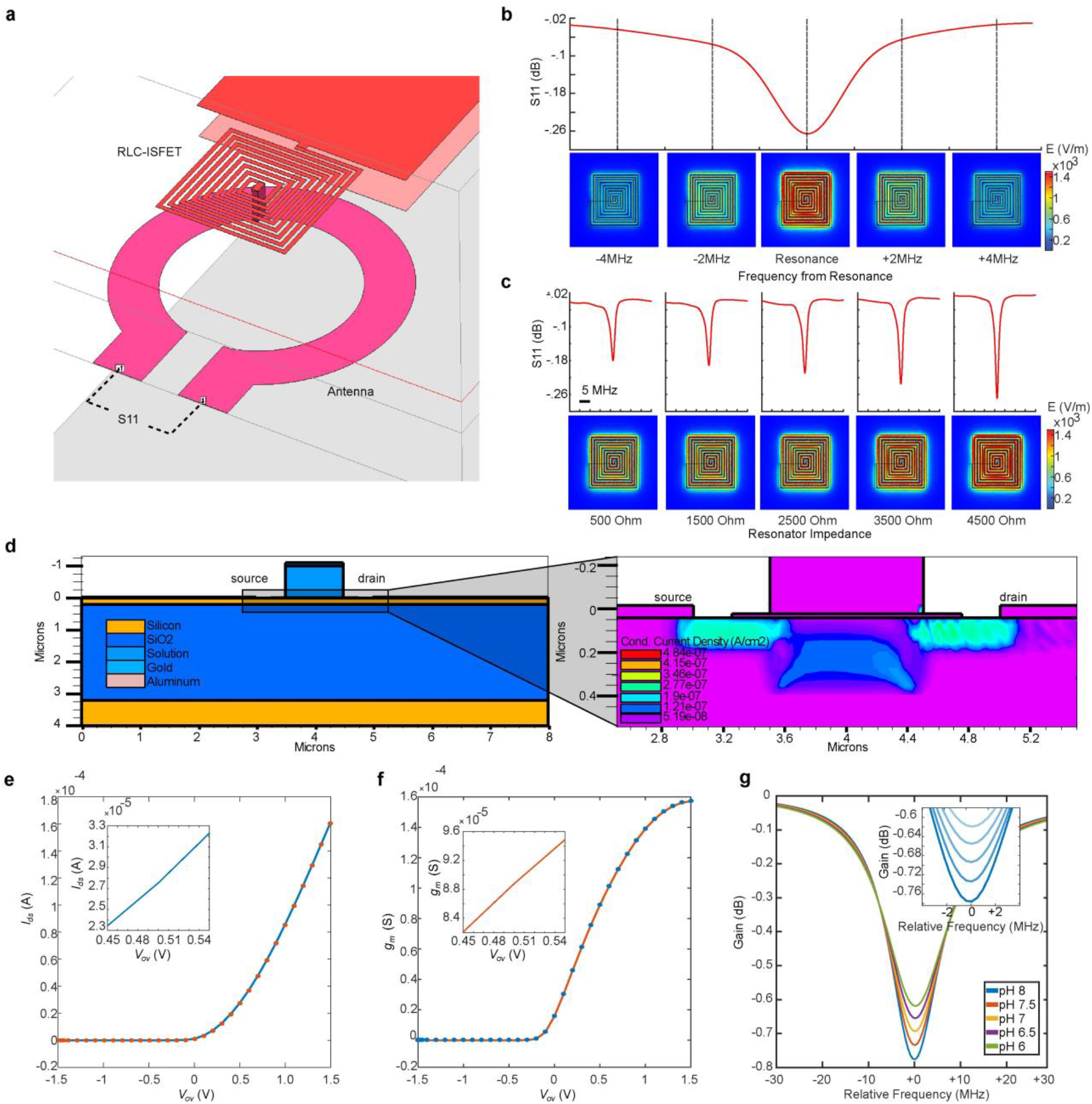
Electromagnetic simulations of ISFET-coupled resonator response. **a** Simulation arena of RLC resonator coupled to ISFET model. S-parameter frequency response is evaluated at a near field receiver. **b** E-field is maximized at resonance. **c** Changes in ionic concentrations at ISFET gate decrease impedance, Q, and e-field. **d** Left - representative ISFET model connected to resonator. Right - current field density at 0.5 V overdrive voltage (*V*_*ov*_). **e** Drain-source current (*I*_*ds*_) as a function of *V*_*ov*_. **f** Small signal transconductance (*g*_*m*_) as a function of *V*_*ov*_. **g** Frequency response modulation at physiological pH range. Inset **-** closeup surrounding resonance.

### 3.2 *In vitro* validation

To test our theoretical predictions and validate the ability of ISFET-coupled resonators to detect physiological pH levels and ion concentrations, we used a simple benchtop configuration to measure response to different pH levels *in vitro* (Fig. 3). Encapsulated ISFET device was bonded to a printed inductor (10 Turns, *d =* 3.5mm) connected in series to a parallel plate capacitor resulting in a resonant frequency of ranging MHz. The device was inductively coupled to a near field antenna (1 Turn, *d =* 9mm) while immersed in test solution for high-speed vector network analyzer measurements (Fig. 3a). For relevant physiological pH levels ranging between 6.5 and 7.5 the test devices displayed Q factor values ranging between 42.3 and 18.1 that decreased by 57.0 % per pH (Fig. 3b). ISFET IV characteristics demonstrated *V*_*th*_ = 0.3 V and *g*_*m*_ = 0.2 mS with a change in *V*_*ds*_ of 55 mV/pH (Fig. 3c). The average RMS noise during acquisition was .0042 dB, corresponding to minimum detectable change of pH = .0229. Detectable concentrations average at 7.57nM. These values correlate well with our theoretical predictions.

**Fig. 3.**
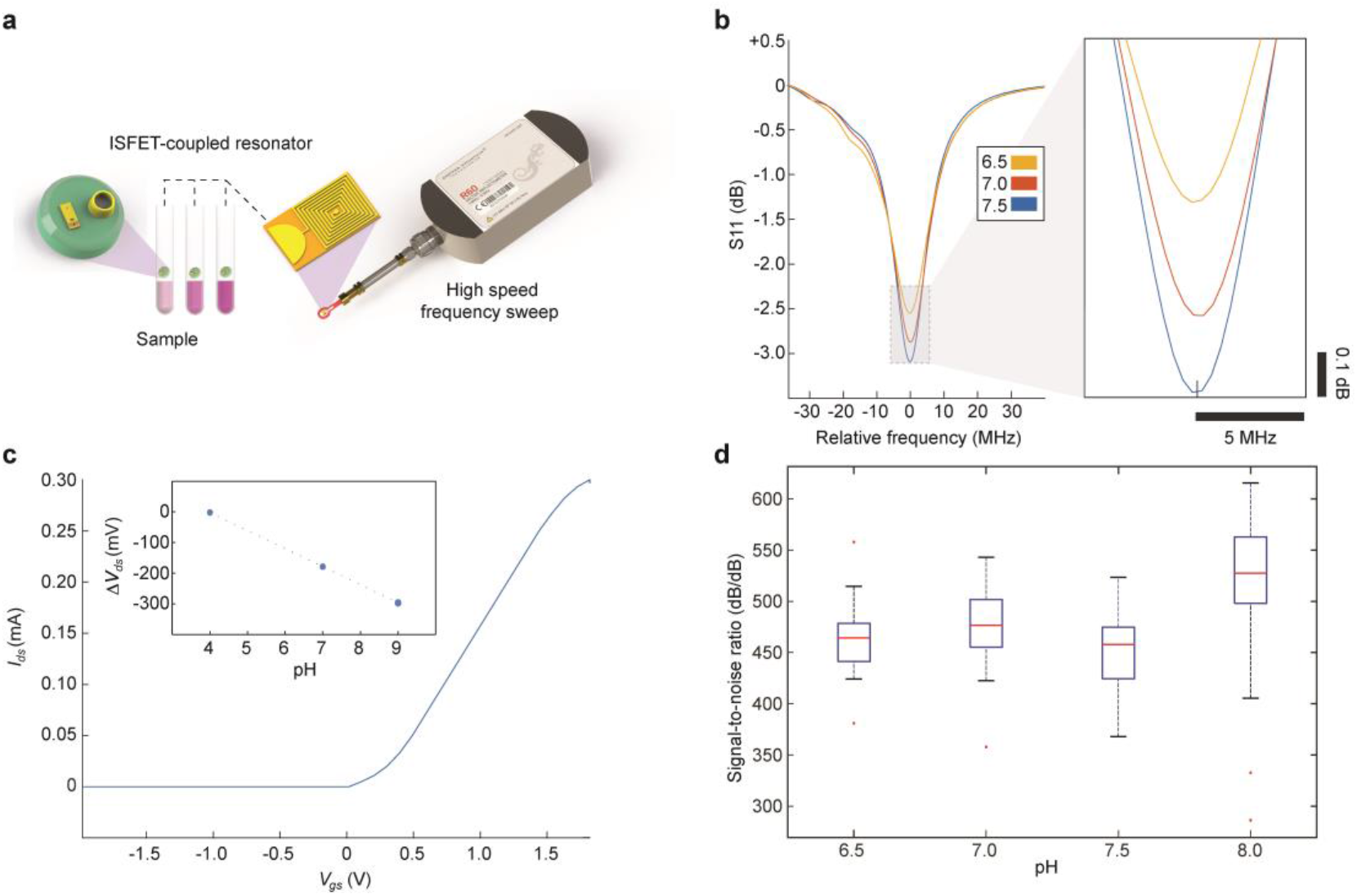
Benchtop assessment of wireless ISFET sensitivity. **a** Experimental configuration: readouts of ISFET-coupled resonator immersed in different pH samples are received by near field antenna. A series of frequency response sweeps is acquired by high-speed vector network analyzer. **b** Examples of frequency response curves for physiological pH levels. **c** Current-voltage (IV) characteristic curve. Inset: Change in drain-source voltage (*V*_*ds*_) with pH levels. **d** Arithmetic mean of signal-to-noise ratio per pH level (red: outliers, included in mean, n = 10, all error bars denote s.e.m.).

### 3.3 *In vivo* recording during hindpaw stimulation

We turned to evaluating the response of ISFET-coupled resonators implanted in the somatosensory cortex (S1HL) of live rats during electrical hindpaw stimulation (Fig. 4). Wireless ISFET (Fig. 4a-c) and wired electrode LFP recordings (4d-e) were acquired during a baseline (no stimulation) period, and during stimulation at frequencies of 2, 5, and 10 Hz. Absolute amplitude of baseline ISFET response was .025 dB, increasing to .08, .04, and .03 dB (*n = 5*) at stimulation frequencies of 2, 5 and 10 Hz, respectively. This corresponded to LFP amplitudes of 0.3 ± 0.25, 1.5 ± 0.75, 1.25 ± 1.0 and 0.8 ± 0.5 mV (*n = 5*) for baseline, 2, 5, and 10 Hz, with largest evoked LFP signal amplitudes observed at 2 Hz and at cortical depths of 1.5 – 1.8 µm (Fig. 4d-e). The ratio between evoked response and baseline amplitudes for all stimulation frequencies was larger than unity (Fig. 4f), with maximal and significant increase of 92.4 ± 44.5 % and 196 ± 73.0 % at 2 Hz stimulation for ISFET and electrode (p < .05 for both, Fig. 4f) and gradually smaller amplitude increases of 24.8 ± 34.3 % and 145 ± 52.1 %, 8.07 ± 28.0 % and 97.4 ± 27.8 % for ISFETand LFP at 5 Hz and 10 Hz. The average normalized LFP response to stimulation degraded linearly for increasingly higher stimulation frequencies and exponentially for ISFET response where average amplitudes for both 5 Hz and 10 Hz stimulation were significantly smaller than 2 Hz (fig. 4d, purple, n = 5, p < .05), but indistinguishable from each other (fig. 4d, magenta, p > .05) and can be attributed to intrinsic nonlinearities of FET operation compared with passive wired electrodes. These values correlate with previous studies demonstrating maximal evoked field potential amplitudes surrounding 2 Hz stimulation of the hindpaw contralateral to the S1HL region of the somatosensory cortex[43].

**Fig. 4.**
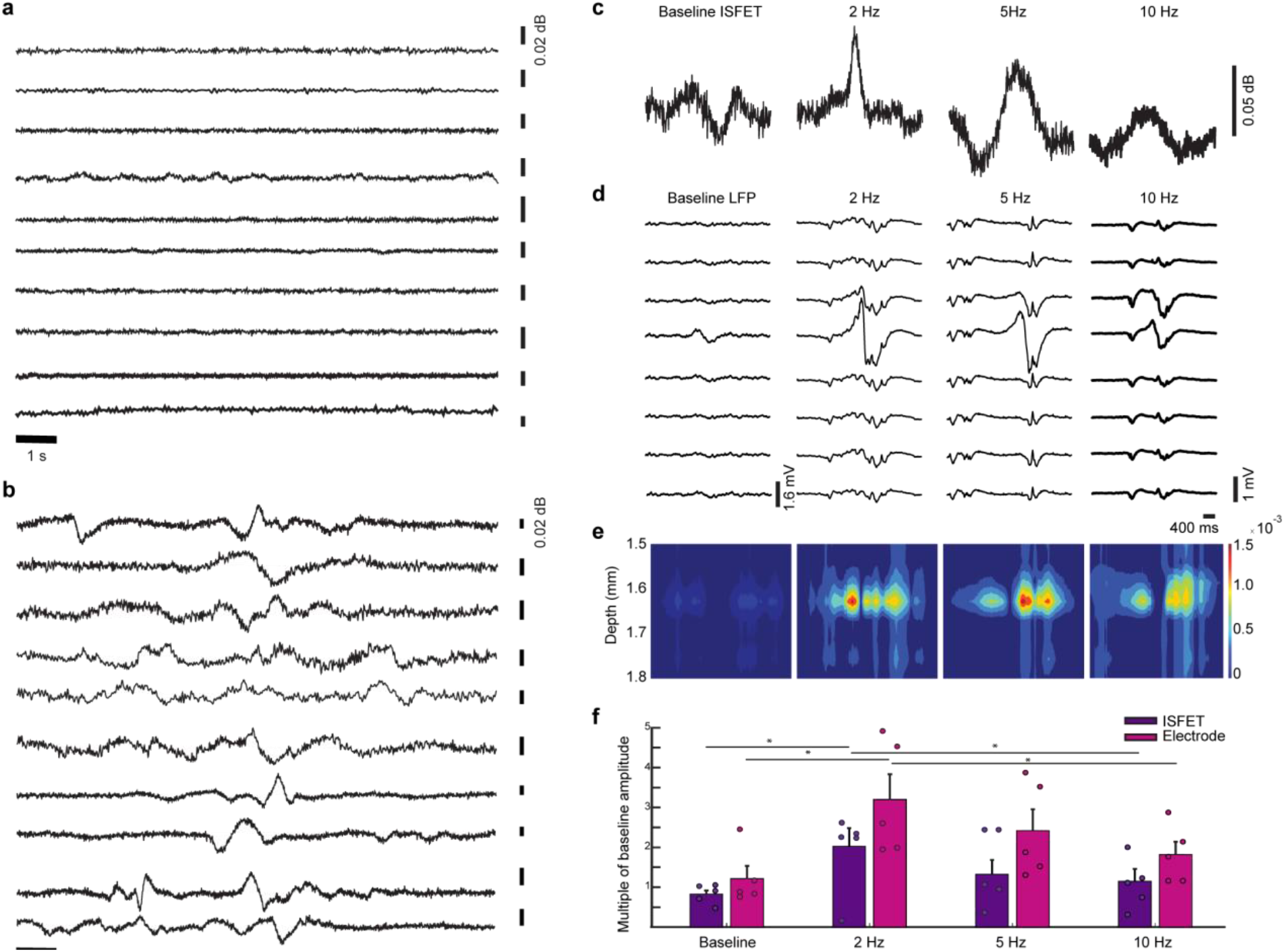
Wireless ISFET somatosensory cortical recordings during hindpaw stimulation. **a** Pre-stimulus readouts of spontaneous activity. **b** Readouts from S1HL somatosensory cortex during a 2 Hz electrical stimulus of contralateral hindpaw. **c** Representative maximum single pulse responses in wireless ISFET recording. **d** Peak differential LFP recordings in response to stimulation. **c** Heatmap depicting maximum peaks of LFP recordings in d. **e** Average amplitude of response to stimulation normalized to baseline. Asterix denotes t-test p-values < .05, error bars are standard errors, n = 5 for all conditions, error bars denote s.e.m.

To identify brain activity bands and temporal signal properties contributing to the recorded signals, we sorted the ISFET activity spikes and performed spectral analysis at frequencies between 0.1 and 8 Hz (Fig. 5). Phasic characteristics of ISFET signals (Fig. 5a-b) spanned a repertoire primarily comprising single phase negative and positive peaks (Fig. 5a, center and left panels) with average duration of 54.0 ± 5.6 ms and 30.8 ± 1.3 ms (Fig. 5b, cyan and yellow points) and positive-phase slow sustained peaks (Fig. 5a, right panels) with average duration of 1.09 ± .128 sec (Fig. 5b, magenta points). Average rise times for each signal type were 10.7 ± .655 ms, 15.1 ± 2.1 ms, and 81.1 ± 15.5 ms for positive, negative, and sustained positive peaks, respectively. Average decay times were 36.6 ± 3.5 ms, 95.5 ± 14.9 ms, and 969.5 ± 122.7 ms for positive, negative, and sustained positive signals, respectively. Power spectrograms of periods following stimulation onset show majority response centered around 0.1 – 5 Hz frequency band and minimal to no response at frequencies larger than 5 Hz (Fig. 5c). Normalized responses for both ISFET spikes and LFPs in the delta band were correlated (Fig. 5d, Pearson correlation r = 0, p = .03, *n = 5*) demonstrating exponential decline in response. As with absolute amplitude (Fig. 4f) no distinct statistical variance could be found between 5 Hz and 10 Hz response in either LFP or ISFET recordings (p > .05). In summary, our recordings demonstrate the ability of the device to record ionic fluctuations correlating ith LFP responses in S1HL following hindpaw stimulation.

**Fig. 5.**
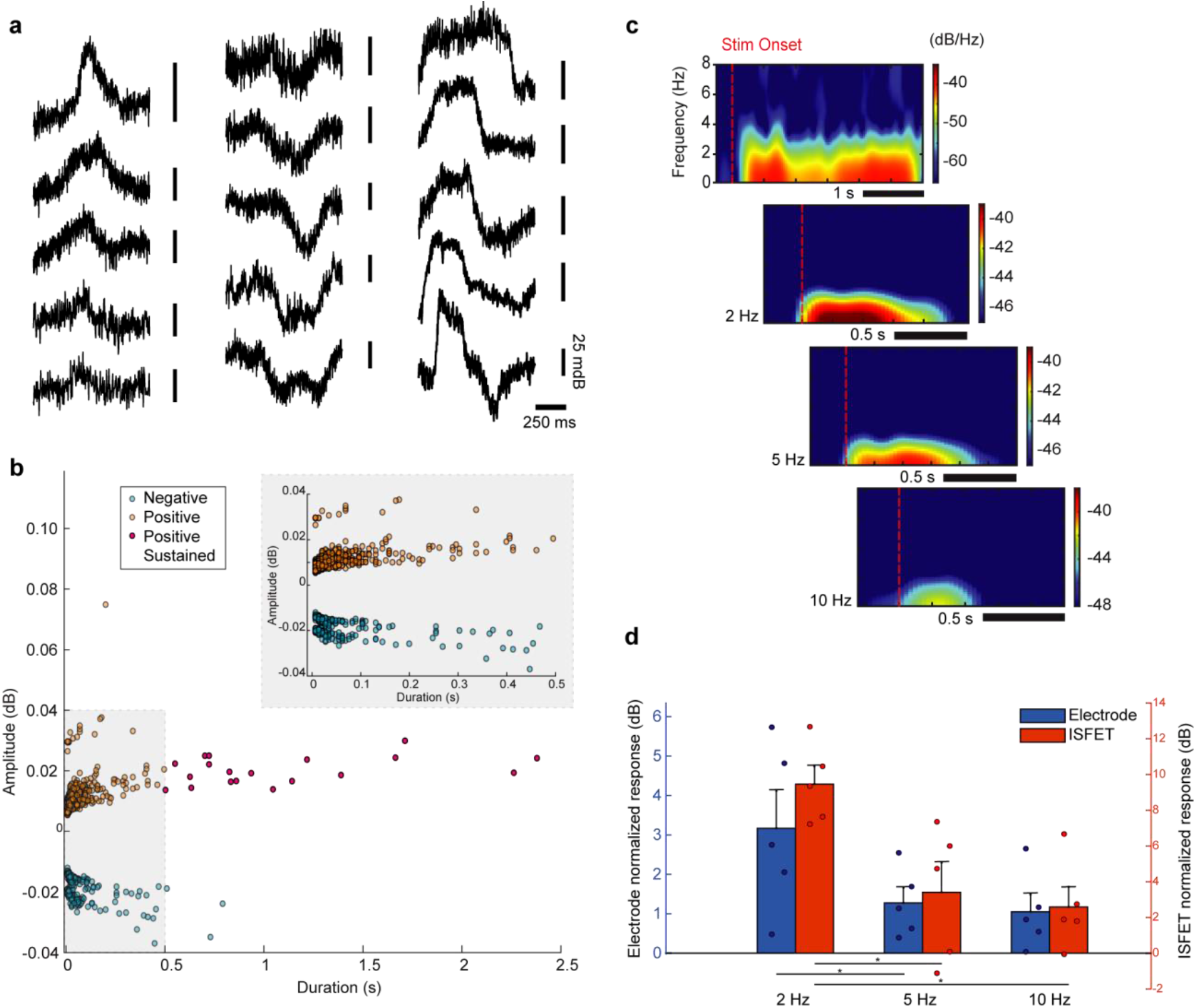
A repertoire of positive and negative phase wireless ISFET responses correlating with delta LFP activity band. **a** ISFET traces can be sorted by both duration and amplitude of response. **b** Majority of ISFET fluctuations are of duration < 500 ms, with positive sustained fluctuations displaying durations of up to 2.5 s. Negative phase responses are of duration < 100 ms. **c** Spectrograms of ISFET wireless response following stimulation onset reveal excitatory response centered around 0.1 - 5 Hz. Duration and intensity of response are inversely proportional to frequency of stimulation. ISFET activity at frequencies greater than the delta wave band was minimal. **d** Response normalized to baseline for both wireless ISFET and LFP electrode recordings show maximal excitatory response at 2Hz stimulation frequency, and a reduced response at frequencies > 5Hz in the delta wave band. Asterix denotes t-test p-values < .05, error bars are standard errors, n = 5 for all conditions, error bars denote s.e.m..

## 4. DISCUSSION

Our results establish a highly compartmentalized wireless technique for *in situ* recording of cortical activity using a standalone ISFET device able to tune an RF resonator in response to neurogenic ionic fluctuations occurring at the ISFET microenvironment. We show that the spatiotemporal characteristics of wirelessly recorded ISFET responses correlate well with LFP recordings tested during standard electrical stimulation protocols of the hindpaw contralateral to the somatosensory cortical implantation site. By utilizing an inductively-powered component serving as a sensitive tunable junction, our approach forms a proof-of-concept for localized transmission of electrophysiological events in the brain to outside antennae or to more sophisticated hardware for neuroimaging modalities operating at the RF regime[44,45,15–17].

Numerous studies showcase the capability of wired transistors to acquire fast millisecond-scale single spikes *in vitro*[29,32,33,46–48] and more recently *in vivo*[49–51]. Alternative transistor designs with optimized geometry and favorable material composition can combine with our tunable sensing architecture and facilitate more sensitive wireless detection of rapid physiological events. Increasing channel transconductance (*g_m_*) beyond ∼1⋅ 10^−4^ S measured within the linear region of the FET on-state for the devices used here is possible using improved designs that rely either on minimization of gate layer thickness[52,53], high performance silicon nanowire gate terminals[54,46,47], polymer-based gate channels for organic electrochemical transistors (OECTs)[53,55] or other ion-gated electrochemical transistors (e-IGTs)[56–58] that display significantly higher mobility and *g_m_* values of up to 1⋅ 10^−2^ S. These and similar devices can serve as more potent tuning switches for *in situ* wireless detection. In addition, the gate channel width of the current device can be modified for optimized *g_m_*, signal-to-noise ratio and threshold voltage[59,60] dictated by the required dimensions of the recording site and critical to overcoming recording artifacts augmented in awake subjects. Incorporating these strategies with the tunable antenna approach can be possible by high speed VNA frequency response acquisition (145 sweeps/sec for 101 discrete frequency measurements in the current study) that could be improved further (>1KHz) via custom hardware and more sophisticated selection of frequency datapoints during sweep acquisition, opening the door to recordings correlated with neurophysiological events manifested at higher frequencies.

Other natural steps for the technology involve integrating the resonator and ISFET components on the same die towards non-surgical injectability into brain parenchyma[19,25,61] similarly to reported RF transmitters used for magnetic sensing and micro-localization *in vivo*[28,62], ingestible electroceuticals for gut therapeutics[63,64], temperature monitoring[65] and wireless neural stimulation[8,66]. A parallel route will be fabrication on mechanically adaptive and biologically-integrated substrates[49,25,61,67,68] to facilitate hermetic chronic operation. Additionally, in order to provide neurochemical specificity, the active site of the ISFET can be functionalized with neurochemical-responsive enzymes[34–36], receptors[69–71] and synthesized entities[72,73] and used to gain greater insight into neurotransmitter levels in the brain, complementing recent efforts demonstrating ISFET-mediated detection of dopamine, serotonin and other neurotransmitters in the brain[36,72,73]. Together, these avenues can lead to biocompatible cellular-scale tunable brain sensors able to sense a milieu of events while mitigating gliosis and related adverse immune responses commonly afflicting chronically implanted recording devices[74,75].

## AUTHOR CONTRIBUTIONS

AH, SB and EM designed the research. SB and EM performed *in vivo* experiments with participation of IB. TZ built and coded the high-speed network analysis interface with JP. SB, JE, MD and AA performed electromagnetic modelling. AV, JG and NG performed finite element simulations. SB and EM analyzed the data. SB, EM and AH wrote the manuscript.

